# Microplastics disrupt adaptive predator-induced behavioural plasticity in *Rana dalmatina* tadpoles

**DOI:** 10.64898/2026.06.11.731648

**Authors:** Dávid Herczeg, Gergely Horváth, Zsanett Mikó, Boglárka Kovács, Attila Hettyey, Gábor Herczeg

## Abstract

The accumulation of microplastics (MP; plastic particles 1 μm – 1 mm in diameter) in the environment is an increasing global concern. Although the physiological effects of MP on organisms, including humans, are increasingly documented, their impacts on behaviour are far less understood. Further, little is known about how MP affect adaptive phenotypic plasticity – the ability of a genotype to adaptively modify its phenotype in response to environmental cues. Anuran tadpoles are key models for studying phenotypic plasticity, with well-established evidence for predator-induced behavioural adjustments. Tadpoles typically reduce their activity and risk-taking when exposed to chemical cues released by predators, which has been proven to be adaptive. We investigated whether MP exposure from the fertilised egg stage alters the behaviour of agile frog (*Rana dalmatina*) tadpoles, and whether it interferes with their predator-induced behavioural plasticity. Tadpoles exposed to chemical cues from dragonfly larvae showed the expected antipredator response: these larvae showed significantly reduced movement activity and risk-taking. Although exposure to MP did not influence the behaviour of tadpoles that had not been exposed to predator cues, and also did not alter predator-induced changes in movement activity, it entirely abolished the predator-induced reduction in risk-taking. These results indicate that MP can compromise antipredator behaviour, increasing the vulnerability of individuals and natural populations to predation without causing visible developmental abnormalities. We recommend that future MP research targets behavioural traits with direct relevance to survival and reproduction, and examines how adaptive phenotypic plasticity is affected.

## Introduction

Humanity has been producing plastics since the 1950s, primarily for broad industrial applications and healthcare (Geyer, 2020). Since then, annual global plastic production has increased exponentially, reaching 430.9 million tonnes in 2024 (2024, 2024; OECD, 2022). In parallel, vast quantities of plastics have been released into terrestrial and aquatic environments, where they accumulate and can persist for centuries (Chamas et al., 2020), raising increasing concern (Horton et al., 2017). According to the recommendation of Hartmann and colleagues (Hartmann et al., 2019), we consider plastic debris to be microplastics (MP) if its particle size falls between 1 mm and 1 μm in diameter. MP exposure can cause physiological and developmental biological malfunctions in a wide array of plant (Li et al., 2022; Shi et al., 2024) and animal taxa (Prokić et al., 2019; Wright et al., 2013); including humans (Marfella et al., 2024; Zolotova et al., 2022). Conversely, changes in behaviour caused by MP (Cozzolino et al., 2025; Horváth et al., 2025; McCormick et al., 2020; Nanninga et al., 2020) or its leachates (Seuront, 2018) have just recently got into the focus of attention. This is surprising because ecologically unjustified behavioural shifts (e.g., changing anti-predator behaviour without a change in predation pressure) can have detrimental effects on survival and reproduction (Lind and Cresswell, 2005). Further, MP can potentially alter adaptive phenotypic plasticity, a genotype’s ability to develop or express different adaptive phenotypes in different environments (West-Eberhard, 2003), again, crucial for survival and reproduction. However, such potential effects are yet scarcely known (Brehm et al., 2026).

Generally, pollution of wetlands occurs *via* the deposition of plastic waste or inappropriate wastewater management (Qian et al., 2021), atmospheric deposition (Allen et al., 2019; Sun et al., 2022), or water draining from agricultural fields (Wang et al., 2022). In this way, MP are carried to surface and sub-surface water bodies, which serve as reservoirs for MP (Di and Wang, 2018; Qian et al., 2021). Accumulating MP in surface water bodies can negatively impact different trophic levels, ultimately affecting interactions in aquatic food webs (Siddiqui et al., 2023; Wang et al., 2019). Amphibian larvae get into contact with MP particles primarily *via* feeding and respiration (Boyero et al., 2020). MP can negatively affect feeding efficiency, growth, development, immune defence, reproductive output, and survival (Araújo et al., 2020; Bosch et al., 2021; Boyero et al., 2020; Cai et al., 2024; Ruthsatz et al., 2022). Larval amphibians present a key model for understanding adaptive behavioural plasticity. For instance, adaptive predator-induced morphological, life-history and behavioural plasticity are common in most studied taxa (Buskirk and Mccollum, 2000; Dijk et al., 2016; Laurila, 2000). Plastic anti-predatory behavioural responses include reduced activity and risk-taking, increased use of cover, spatial avoidance, and enhanced shoaling behaviour (Relyea, 2003; Sergio et al., 2021; Skelly and Werner, 1990; Urszán et al., 2018; Watt et al., 1997). However, the effects of MP exposure on these behaviours remain poorly studied in amphibians.

The present study had two aims: to see whether permanent MP exposure, starting from the fertilised egg stage, (i) affects movement activity and risk-taking of anuran larvae (tadpoles) and (ii) interferes with adaptive behavioural plasticity induced by perceived risk of predation. Our model was the widespread European agile frog, *Rana dalmatina* (Bonaparte 1840), where the occurrence of clear predator-induced behavioural plasticity has been documented (Hettyey et al., 2011; Hettyey et al., 2010; Urszán et al., 2015). We employed microparticle treatments (control, silica powder (SiO_2_) and MP) and predatory treatments (presence/absence of chemical cues from a common predator, *Aeshna cyanea* dragonfly larvae) in a factorial design. We hypothesized that MP exposure impacts both movement activity and risk-taking behaviour of the tadpoles *per se*, and also interferes with their predator-induced behavioural plasticity, leading to weakened antipredator responses. The predicted outcome would imply that tadpoles exposed to MP may become more vulnerable to predation in their natural habitat, resulting in increased mortality and potential population declines in the wild.

## Materials and Methods

All experimental procedures were approved by the Ethical Commission of the Plant Protection Institute, HUN-REN ATK, and permissions were issued by the Government Agency of Pest County (PE/EA/00270-6/2023). The experiments were carried out according to recommendations of the EC Directive 86/609/EEC for animal experiments (http://europa.eu.int/scadplus/leg/en/s23000.htm). We used exclusively metal or glass equipment during the investigation except in the case when tadpoles were kept individually in opaque white polypropylene plastic boxes to minimise the risk of cross-contamination with MP. Following each water change, wastewater was pre-filtered prior to disposal.

### Animal husbandry and pre-hatching treatment

In March 2025 we collected 10 freshly laid egg clutches of *R. dalmatina* from a pond in a forested area of the Visegrádi hills’ pond (47°46’01.4"N, 18°58’54.0"E) and transported them to the laboratory at the Plant Protection Institute, HUN-REN ATK on the outskirts of Budapest. We assessed background microplastic contamination in the pond (covering particles within the 50–1000 μm size range) using FTIR spectroscopy and detected only very low polymer concentrations, with a total concentration of 0.024 particles L⁻¹ and a polystyrene concentration of 0.006 particles L⁻¹ (Herczeg D., pers. comm.). On Day 0, we separated 9 partial clutches into three replicates, each containing approx. 100 eggs (103.6 ± 13.5, mean ± SD) and reared them in glass aquaria (20 cm × 13 cm × 22 cm, length, width, height, respectively) holding 0.5 litre of reconstituted soft water (RSW; 48 mg NaHCO_3_, 30 mg CaSO_4_ × 2 H2O, 61 mg MgSO_4_ × 7 H_2_O, 2 mg KCl added to 1 liter reverse-osmosis filtered, UV-sterilized tap water). One clutch was infertile and was therefore excluded from the experiment. The three replicates of each clutch were randomly assigned to one of the following pre-hatching microparticle treatment groups: (i) control, with no microparticles added; (ii) natural microparticle, with silica powder (SiO₂) added; and (iii) MP, with polystyrene added to the rearing water of animals. We used pristine, spherical polystyrene MP (Thermo Fisher Scientific™, Fluoro-Max™ Fluorescent beads, Cat. No. CDG1000) with a particle size of 10 µm, which are known to be consumed by tadpoles (Araújo et al., 2020; Bosch et al., 2021; Boyero et al., 2020). Polystyrene was chosen due to its high prevalence in freshwater environments (Jones et al., 2020; Wagner et al., 2014). We used SiO₂ powder (mean particle size 17 μm; Sigma-Aldrich, product number: 227196) as a natural microparticle (Kotta et al., 2022) to mimic diatoms, which are commonly ingested by tadpoles (Ranvestel et al., 2004). We applied a relatively high concentration of MP (180 particles/ml), still falling within the environmentally relevant range of surface-water MP contamination (Koelmans et al., 2019) and has been used in previous studies on tadpoles (Araújo et al., 2020; Bosch et al., 2021; Boyero et al., 2020; Martin et al., 2025; Ruthsatz et al., 2023). The same concentration was used in the SiO₂ treatment groups. In the control group, we added an equivalent volume of RSW to the rearing water of tadpoles.

### Experimental design

We captured 15 dragonfly larvae (*Aeshna cyenea*) from a nearby artificial pond (47°33’04"N, 18°55’36"E) and transported them to the laboratory. We placed predators individually in 500 mL cups (H 13.5 cm; radius 9 cm) holding 300 mL RSW and a birchwood stick (20 cm long and 0.4 cm in diameter) as a perching site and fed them with two naïve agile frog tadpoles originating from the same habitat every other day. We assigned predators to three groups, designed to obtain all types of chemical cues both from predators as well as from their prey (Hettyey et al., 2015): (i) predators fed two naïve tadpoles in the morning, (ii) predators fed two naïve tadpoles in the afternoon (shortly before predator treatment), and (iii) predators not fed tadpoles, i.e., starved predators (Van Buskirk and Arioli, 2002). The feeding regime of predator groups was alternated daily. On Day 16, when the majority of tadpoles had fully developed oral discs and started feeding independently, while the operculum was formed and external gills disappeared (developmental stage 25, (Gosner, 1960)), we haphazardly selected 12 healthy-looking individuals from each aquarium and kept them from here on individually in boxes (L 24 × W 16 × H 13 cm) in 1 L RSW. We started the 3 × 2 factorial experiment with the same three levels of microparticle treatments as applied during tadpole rearing (control, SiO_2_, and MP), and employed two predator-cue treatments. Predator cues from the three groups were pooled and 5 ml were added with glass syringe to each tadpole during the afternoons from Day 19, five times per week until the end of the experiment. Non-predator treatment groups received an equivalent volume of RSW. We changed the RSW in the rearing boxes twice per week while maintaining the initial microparticle and predator cue treatment concentrations. Following each water change, we fed tadpoles *ad libitum* with chopped, slightly boiled spinach. Tadpoles not used in the experiment were transported back to the collection site.

On Day 30, two days after the last water change, the same observer (DH) recorded tadpole behaviour by eye at hourly intervals between 10:00 and 14:00 (five observations), using a telescopic mirror to avoid disturbing the animals. Two behaviours were assessed: movement activity (actively moving tadpole and/or moving tail of tadpole *vs*. unmoved tadpole) and risk-taking (being on the bottom, representing risk-averse, or in the water column, representing risk-prone behaviour).

### Statistical analysis

To test treatment effects on activity and risk-taking, we fitted generalized linear mixed models (GZLMMs) with a binomial distribution using the *lme4* (Bates et al., 2015) and *lmerTest* (Kuznetsova and Christensen, 2017) packages in R version 4.5.1 (R Core Team, 2025). Activity and risk-taking were positively correlated (Pearson’s *r* = 0.49, 95% confidence interval [CI]: 0.45 – 0.53, *P* < 0.001). Nevertheless, movement activity and risk-taking represent biologically distinct behaviours (Réale et al., 2007). Therefore, we fitted separate GZLMMs for both behavioural traits, specifying the focal behaviour as a binary response variable in each model. To account for the observed correlation, we included each behaviour as a fixed effect in the model of the other: tadpole position (bottom *vs.* water column) was included as a fixed effect in the activity model, whereas tadpole activity (mobile *vs*. immobile) was included as a fixed effect in the risk-taking model. Predator-cue treatment (present *vs.* absent) and microparticle treatment (control *vs.* SiO_2_ *vs.* MP) were included as fixed effects, along with the two-way interactions and the three-way interaction among the fixed effects. To control for within-day habituation, we added the centered (mean = 0, SD = 1) order of observations to the models. Individual identity was added to models as a random effect. Fixed effects were tested using Wald’s chi-squared tests and random effects by likelihood ratio tests. *P*-values for the likelihood ratio tests were calculated following Zuur et al. (Zuur et al., 2009). The model’s estimated marginal means were extracted using the *emmeans* package (Lenth et al., 2021). To compare groups, we looked for the presence/absence of overlaps between 85% CIs following Payton et al. (Payton et al., 2003), who demonstrated that the lack of overlap in 83–84% CIs is analogous to a *P*-value < 0.05.

## Results

Tadpole activity was significantly affected by the predator-cue treatment (Table 1): the number of active tadpoles in the presence of predator cues was substantially reduced of that observed in the absence of predator cues (Fig. 1a). On the other hand, we did not find an effect of microparticles, or predator × microparticle interaction (Table 1). Tadpoles in the water column were more likely to be active than those on the bottom, while the microparticle × position interaction was also significant (Table 1): tadpoles exposed to MP were less active on the bottom of the box compared to SiO_2_-exposed conspecifics (odds ratio: 0.602 ± 1.145 SE, z. ratio: -2.094, *P* = 0.049). In contrast, tadpoles exposed to MP were significantly more active in the water column, compared to the SiO_2_-exposed conspecifics (odds ratio: 1.665 ± 0.404 SE, z. ratio: 2.099, *P* = 0.049). Remaining non-significant effects are shown in Table 1.

**Figure 1.**
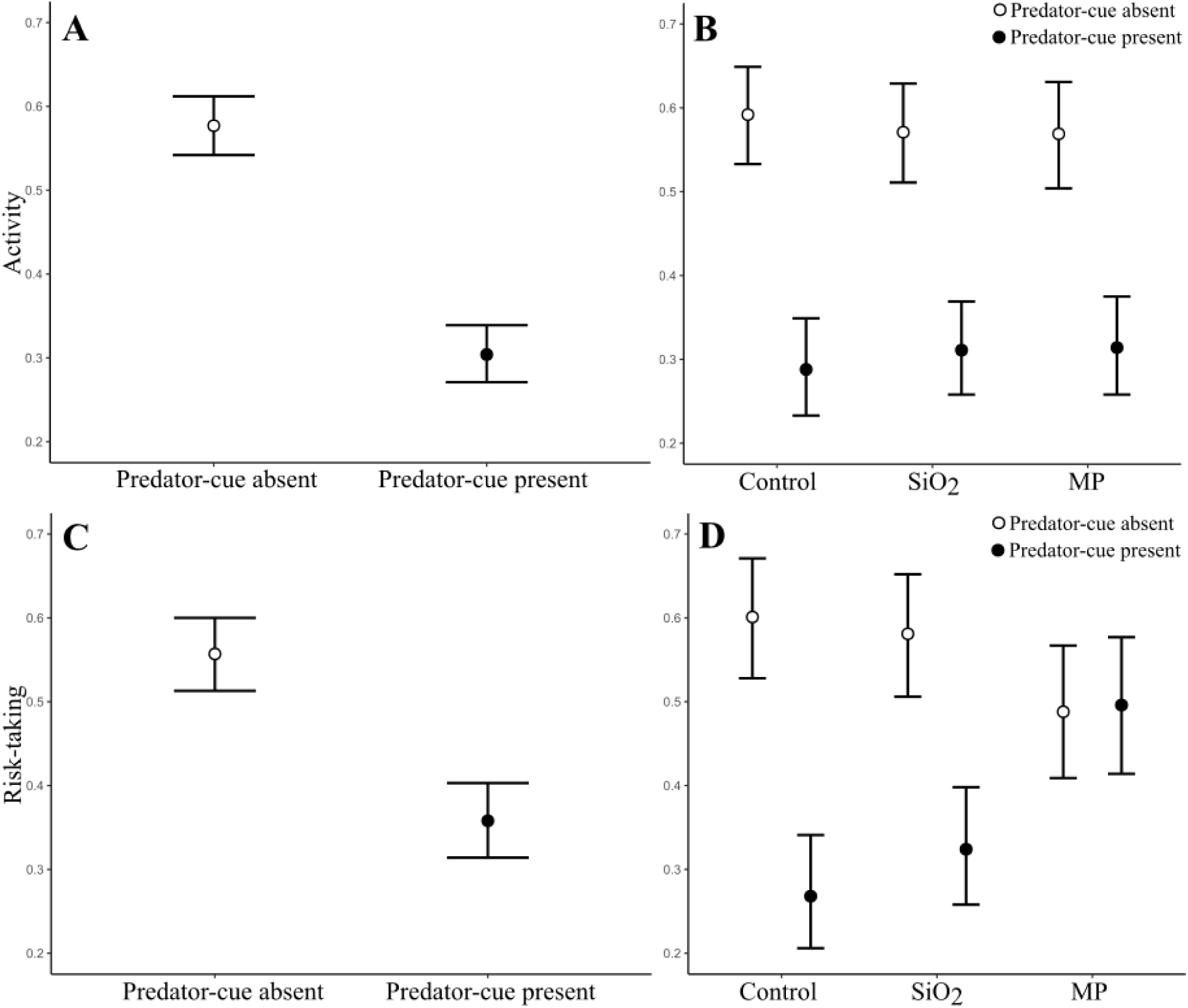
Effect of predator-cue absence versus presence on the activity and risk-taking behaviour of *Rana dalmatina* tadpoles. Activity is shown for (A) the pooled sample and (B) across different microparticle treatments. Risk-taking behaviour is presented in the same way (C, D). Points represent estimated marginal means, and error bars indicate 85% confidence intervals (CIs). Non-overlapping 85% CIs indicate a statistically significant difference. (Payton et al., 2003). Control: no-microparticle; SiO_2_: silica powder; MP: microplastics

**Table 1.** The ANOVA table of GZLMMs on activity and risk-taking of *Rana dalmatina* tadpoles. Significant effects are in bold. Correction indicates the correction for the position in the case of activity, while correction for activity in the case of risk-taking.

| **Model term** | **Activity** | **Risk-taking** |
| --- | --- | --- |
| | $\chi^2$ (df), *P* | $\chi^2$ (df), *P* |
| *Fixed effects* |  |  |
| **Intercept** | **12.21 (1), <0.001** | **3.87 (1), 0.049** |
| **Predator-cue (Pr)** | **57.32 (1), <0.001** | **19.77 (1), <0.001** |
| **Microparticle** | <0.01 (2), 0.99 | 1.42 (2), 0.49 |
| **Order of tests** | 0.26 (1), 0.61 | 0.71 (1), 0.39 |
| **Correction (Corr)** | **231.09 (1), <0.001** | **218.37 (1), <0.001** |
| **Pr × Microparticle** | 0.45 (2), 0.79 | **11.23 (2), 0.003** |
| **Pr × Corr** | 0.27 (1), 0.6 | 0.09 (1), 0.76 |
| **Microparticle × Corr** | **9.76 (2), 0.007** | **11.24 (2), 0.003** |
| **Pr × Microparticle × Corr** | 1.94 (2), 0.38 | 1.86 (2), 0.39 |
| *Random effect* |  |  |
| **Individual** | **12.32 (1), <0.001** | **55.15 (1), <0.001** |

Similarly, relative tadpole risk-taking was significantly affected by the predator-cue treatment (Table 1): tadpoles were more risk-averse in the presence of predator cues (Fig. 1b). There was no effect of microparticles, however, the predator × microparticle interaction did have a significant effect on risk-taking (Table 1): in the presence of MP, the predator-induced decrease in risk-taking vanished (Fig. 2b). Active tadpoles were more likely to be in the water column than inactive tadpoles Tadpoles in the water column were more likely to be active than those on the bottom and the microparticle × activity interaction was significant (Table 1): active tadpoles were more risk-prone in the MP-exposed group compared to the microparticle-unexposed (odds ratio: 0.524 ± 0.16 SE, z. ratio: -2.122, *P* = 0.046) and the SiO_2_-exposed (odds ratio: 2.163 ± 0.643 SE, z. ratio: 2.597, *P* = 0.014) groups, but in inactive tadpoles, there was no difference in risk-taking between microparticle treatments (all *P* > 0.143). Remaining non-significant effects are shown in Table 1.

## Discussion

Microplastics are pollutants found in literally all habitats on Earth. While knowledge on direct negative effects of MP on survival and development is accumulating at a fast pace in various organisms, information on more subtle effects on seemingly healthy individuals, like changes in behavioural strategies or in the expression of phenotypic plasticity therein is lacking. Under predation pressure tadpoles must adjust their phenotypes to increase their chances of survival. The first line of defence against predators is behavioural plasticity, which is well-documented in tadpoles (Buskirk and Mccollum, 2000; Dijk et al., 2016; Laurila, 2000; Van Buskirk and Arioli, 2002). In the present experiment, our most salient findings are that exposure to MP did not have a direct effect on behaviour *per se*, but it did completely erase the predator-induced decrease in risk-taking.

Studies examining MP-induced changes in anti-predator behaviour vary widely in polymer type, particle shape, concentration, and exposure duration. Despite this heterogeneity, a growing body of evidence suggests that MP or their leachates can directly affect behaviours that are crucial for anti-predatory responses in many organisms (Araújo and Malafaia, 2020; Hawke et al., 2024; Horváth et al., 2025; McCormick et al., 2020; Nanninga et al., 2020; Seuront, 2018; Wang et al., 2023), thereby indirectly impairing phenotypic plasticity. Activity represents a key behavioural trait closely associated with predation risk (Wellborn et al., 1996). Increased activity boosts both food intake and susceptibility to predation (Anholt and Werner, 1995; Skelly, 1994). Here we found decreased activity under perceived predation risk but detected no change in the activity of tadpoles following prolonged MP exposure. Risk-taking behaviour has not previously been examined in studies investigating the anti-predator behavioural effects of microplastic exposure in amphibians (Araújo and Malafaia, 2020; Scribano et al., 2023). As expected, agile frog tadpoles were more risk-averse in the presence of predator cues, but again, we found no direct effects of MP on risk-taking.

However, not only direct effects of MP on behaviour can make the affected individuals vulnerable to predation. If MP abolish naturally occurring predator-induced adaptive behavioural plasticity, individuals may behave maladaptively, and the lack of a response to predator presence may result in high mortality risk. The anti-predator response of decreased activity was the same in all MP treatments. Similarly, in a recent study, tadpoles of the Italian agile frog *Rana latastei* reduced their activity in the presence of predator cues from dragonfly larvae, whereas exposure to different concentrations (1, 7, and 50 mg L^−1^) of a mixture of plastic polymers (HDPE, PVC, PS and PES) for two weeks did not significantly alter the strength of this behavioural response (Scribano et al., 2023). In our study, however, the anti-predatory decrease in risk-taking was similar in the control and SiO_2_ treatments, but it completely vanished in tadpoles exposed to MP.

Microplastics-induced disruption of predator-induced behavioural plasticity may increase the vulnerability and mortality of individuals in natural populations. Such effects might contribute to population declines and challenge ecosystem resilience without visually detectable effects on organismal development. We urge testing MP effects on other ecologically relevant behaviours other than movement activity, such as risk-taking, exploration, sociability and aggression. Further, we recommend elevating the focus from direct effects on behaviour to potential effects on adaptive behavioural plasticity induced by natural stressors, like predation, competition or parasites/pathogens, preferably in a more natural setup than the one employed in our work. Such setups might be based on seminatural enclosures or mesocosms, and MP exposure should include environmentally derived, irregularly shaped MP of various kinds to enhance the ecological relevance of results. In conclusion, simply studying direct physiological or developmental effects is not sufficient for fully perceiving the negative effects of MP on individuals, populations or ecosystems.

## Acknowledgement

We thank Enikő Kovács, Márk Szederkényi, Andrea Kásler, János Ujszegi and Norbert Vörös for technical assistance during the experiment. We thank Fruzsina Gresina (HUN-REN CSFK) for the physical characterisation of SiO_2_ powder for the experiment. Project no. [STARTING 149841 for DH] and [K-147500 for AH] has been implemented with the support provided by the Ministry of Culture and Innovation of Hungary from the National Research, Development and Innovation Fund, financed under the [National Research Excellence Program STARTING_24 subprogram, and Thematic Research Projects Programme 2023] funding scheme. University Excellence Scholarship Program (EKÖP-24-4 to ZM) of the Ministry for Culture and Innovation from the source of the National Research, Development and Innovation Fund. DH and GeH were supported by the János Bolyai Research Scholarship of the Hungarian Academy of Sciences. This project has received funding from the HUN-REN Hungarian Research Network.

## Data statement

The dataset analysed during the current study are available in the Figshare Digital Repository (DOI: 10.6084/m9.figshare.32620290). The DOI becomes active upon acceptance of the manuscript. For reviewers please see the following private link to get access to the data and code: https://figshare.com/s/07f135c90ee9eeebb3b2

## Authors’ contributions

DH and GáH conceived the study and designed the methodology; DH, ZM, BK, and AH collected the data; GeH analysed the data; DH and GáH led the writing of the manuscript. All authors read and approved the final manuscript

